# OncoMORPHIA: An Integrated Web Platform for Interactive 3D Visualization and Functional Annotation of Cancer Mutations

**DOI:** 10.64898/2026.04.01.715816

**Authors:** Mladen Cimeša, Aleksandra Sokić

## Abstract

**Summary:** The rapid accumulation of cancer genomic data across repositories such as ClinVar, cBioPortal, and the TCGA Pan-Cancer Atlas has created an urgent need for integrated tools that allow researchers to explore mutations in their structural and clinical context without requiring specialized bioinformatics expertise. Here we present OncoMORPHIA, a free, browser-based web platform that unifies 3D protein structure visualization, clinical variant annotation, drug-target interaction mapping, survival analysis, mutational signature decomposition, and AI-powered interpretation within a single interface. OncoMORPHIA automatically retrieves and integrates data from ten public databases, maps missense mutations onto experimentally determined or AlphaFold-predicted protein structures, computes mutation density heatmaps with Gaussian smoothing, and renders interactive visualizations including lollipop plots, Kaplan-Meier survival curves, protein-protein interaction networks, and pan-cancer tissue distribution charts. The platform supports 45 major cancer driver genes with extensibility to any human gene with available structural data.

**Availability:** OncoMORPHIA is freely available at https://oncomorphia.com. Source code is available upon request. The platform requires no installation, no account registration, and no API keys for core functionality.

## 1. Introduction

The characterization of somatic mutations in cancer has been transformed by large-scale sequencing initiatives including The Cancer Genome Atlas (TCGA), the International Cancer Genome Consortium (ICGC), and clinical sequencing programs such as MSK-IMPACT. These efforts have catalogued millions of variants across hundreds of cancer types, generating a wealth of data that is distributed across multiple specialized databases: ClinVar for germline pathogenicity classifications, cBioPortal for somatic mutation profiles and clinical outcomes, the Protein Data Bank (PDB) for experimentally determined structures, and AlphaFold for computationally predicted models.

A central challenge facing cancer researchers and clinical geneticists is the fragmentation of these resources. To assess the functional significance of a specific mutation, a researcher must typically navigate between five or more web interfaces: querying ClinVar for pathogenicity, cBioPortal for somatic frequency, the PDB for structural context, DGIdb or ChEMBL for drug interactions, and ClinicalTrials.gov for active trials. This workflow is time-consuming, error-prone, and inaccessible to researchers without bioinformatics training. Furthermore, the January 2024 restructuring of ClinVar’s classification system—which split the single clinical_significance field into separate germline_classification, oncogenicity_classification, and clinical_impact_classification fields—has broken many existing automated pipelines that relied on the legacy format.

Several tools have been developed to address aspects of this challenge. cBioPortal provides comprehensive genomic data exploration but limited structural visualization. COSMIC offers mutation catalogues with lollipop plots but no 3D structural mapping. The RCSB PDB viewer provides excellent structural visualization but no mutation overlay capability. ProteinPaint and MutationMapper offer interactive lollipop plots but lack integration with drug databases, survival data, or AI-powered interpretation. No existing tool combines all of these capabilities in a single, freely accessible, browser-based platform.

Here we present OncoMORPHIA (Oncology Mutation and Residue Phenotype Hub for Interactive Analysis), a web application that integrates data from ten public databases into a unified analytical environment. OncoMORPHIA automatically retrieves mutations, maps them onto 3D protein structures, overlays drug-target interactions, computes survival correlations, and provides AI-powered variant interpretation—all within a single browser session requiring no installation or specialized software.

## 2. Implementation

### 2.1 Architecture and Technology Stack

OncoMORPHIA is implemented as a Python web application using the Streamlit framework (version 1.30+), chosen for its ability to deliver interactive data applications with rapid development cycles. The application is deployed on Railway (railway.app) for production hosting with persistent containers, automatic SSL provisioning, and environment-based configuration management. The 3D molecular viewer is rendered using py3Dmol, a Python wrapper around the 3Dmol.js WebGL library, embedded within the Streamlit interface via HTML components. Interactive lollipop plots are rendered using Plotly for client-side interactivity with hover tooltips, zoom, and pan capabilities. Static visualizations including domain tracks, survival curves, co-occurrence heatmaps, and mutational signature spectra are generated using Matplotlib.

### 2.2 Data Integration Pipeline

OncoMORPHIA integrates data from the following sources through their respective REST APIs, with all requests cached using Streamlit’s built-in TTL cache to minimize redundant network calls:

#### ClinVar (NCBI E-utilities)

Germline variant classifications retrieved via eSummary JSON API. The parser handles both the legacy clinical_significance field and the post-January 2024 germline_classification, oncogenicity_classification, and clinical_impact_classification fields, ensuring compatibility with the current ClinVar data model. Pathogenicity tier resolution uses longest-match-first substring matching to prevent misclassification of composite terms (e.g., “likely pathogenic” matching before “pathogenic”).

#### cBioPortal (REST API v2)

Somatic mutation profiles from MSK-IMPACT 2017 (10,945 tumour samples) and the TCGA Pan-Cancer Atlas 2018 (32 individual cancer-type studies, >10,000 patients). Mutations are queried via the POST /mutations/fetch endpoint using molecularProfileIds arrays for cross-study queries. Survival data (OS_MONTHS, OS_STATUS) are retrieved as patient-level clinical attributes. Pan-cancer tissue distribution is derived from study identifiers, with each of the 32 TCGA studies mapped to its cancer type abbreviation.

#### RCSB PDB and AlphaFold

Protein structures are selected using a mutation-coverage scoring algorithm. For each gene, up to 20 RCSB PDB entries are evaluated by computing the intersection of mutated residue positions with resolved C-alpha atoms. The structure maximizing coverage at the best resolution is selected. When RCSB coverage falls below 50% of mutation positions, the pipeline falls back to AlphaFold predicted structures (v2–v4), which provide full-length models with per-residue confidence scores (pLDDT) embedded in the B-factor column.

#### DGIdb and ChEMBL

Drug-gene interactions are queried from DGIdb (Drug Gene Interaction Database) and cross-referenced with ChEMBL mechanism-of-action and maximum clinical phase data. Results are deduplicated by drug name and sorted by clinical development phase.

#### STRING DB

Protein-protein interaction networks are retrieved from STRING (Search Tool for the Retrieval of Interacting Genes/Proteins) with a minimum combined score threshold of 0.4. Interaction partners are annotated as known cancer genes when present in the curated Entrez ID table, and functional enrichment terms are retrieved for pathway context.

#### Ensembl VEP

Variant effect predictions (SIFT, PolyPhen-2, CADD) are obtained from the Ensembl Variant Effect Predictor REST API via batch POST requests, with rate limiting to comply with API usage guidelines.

#### ClinicalTrials.gov (API v2)

Active recruiting trials matching gene-mutation-cancer queries are retrieved with NCT identifiers, phase information, and intervention details.

### 2.3 Visualization Modules

#### 3D Structure Viewer

The molecular viewer supports four rendering modes: (1) Cartoon mode with pathogenicity-coloured mutation spheres (red = pathogenic, orange = likely pathogenic, yellow = VUS, purple = somatic, green = benign); (2) Spheres mode with stick representations; (3) Heatmap mode projecting Gaussian-smoothed mutation density onto the molecular surface with a configurable smoothing parameter (σ = 0–10); and (4) pLDDT mode (AlphaFold structures only) colouring the backbone by prediction confidence using the standard AlphaFold colour scheme (dark blue ≥ 90, light blue 70–90, yellow 50–70, orange < 50). Mutation spheres are interactive: hover displays mutation name, pathogenicity, and count; click zooms to the residue.

#### Interactive Lollipop Plot

Mutation frequency is visualized as a Plotly-based lollipop plot with vertical stems at each mutated position and circles sized by count and coloured by pathogenicity. Hover tooltips display residue position, mutation count, classification, and specific mutation names. A protein domain track drawn from UniProt annotations is rendered below the frequency axis, providing structural context for mutation clustering within functional domains.

#### Kaplan-Meier Survival Analysis

Overall survival curves are computed using a custom Kaplan-Meier estimator comparing patients with mutations in the query gene (across all 32 TCGA Pan-Cancer Atlas studies) against wild-type patients. Confidence intervals are displayed as shaded bands. Median survival times are annotated on the plot. A companion bar chart shows mutation frequency by residue position to link survival differences to specific hotspots.

#### Mutational Signatures

The six-channel substitution spectrum (C>A, C>G, C>T, T>A, T>C, T>G) is computed from TCGA Pan-Cancer mutation data by normalising reference alleles to pyrimidine context. The spectrum is rendered using COSMIC standard colours and accompanied by a curated lookup table of known COSMIC mutational signature associations for major cancer genes, linking observed mutation patterns to their likely mutagenic aetiologies (e.g., UV exposure, APOBEC activity, mismatch repair deficiency, tobacco smoking).

### 2.4 AI-Powered Mutation Interpretation

OncoMORPHIA incorporates a context-aware AI assistant powered by the Groq inference platform running LLaMA 3.3 70B. The assistant receives the full loaded mutation dataset as structured context in its system prompt, including total mutation count, pathogenicity distribution, top hotspot positions with counts, and individual mutation entries with HGVS notation and clinical significance. This context injection ensures that AI responses are grounded in the actual loaded data rather than relying solely on the model’s training knowledge. Pre-configured quick-question buttons provide one-click access to common queries about hotspot biology, pathogenicity, therapeutic targets, and gene function.

## 3. Features and Functionality

OncoMORPHIA provides thirteen analytical modules accessible via a tabbed interface:

**(1) 3D Viewer** — interactive protein structure visualization with four rendering modes and mutation overlay; **(2) Gene Comparison** — side-by-side analysis of two cancer genes with synchronized viewers and summary statistics; **(3) Mutation Table** — filterable, downloadable table of all retrieved mutations with pathogenicity annotations; **(4) Lollipop Plot** — interactive mutation frequency visualization with domain overlay; **(5) Drugs & Trials** — drug interactions from DGIdb/ChEMBL with clinical trial listings from ClinicalTrials.gov; **(6) Domain Track** — linear protein domain diagram from UniProt with mutation density histogram; **(7) VEP Predictions** — SIFT, PolyPhen-2, and CADD scores with a concordance scatter plot; **(8) PPI Network** — STRING DB interaction partners with radial network graph and functional enrichment; **(9) Co-occurrence** — mutation co-occurrence heatmap from per-sample MSK-IMPACT or TCGA data; **(10) Survival Analysis** — Kaplan-Meier curves from TCGA Pan-Cancer Atlas with pan-cancer tissue breakdown; **(11) Mutational Signatures** — six-channel substitution spectrum with COSMIC signature associations; **(12) Pathway View** — multi-gene pathway diagram with mutation burden sizing across six predefined cancer pathways; **(13) AI Assistant** — context-aware natural language interface for mutation interpretation.

Upon gene selection and analysis initiation, OncoMORPHIA executes a comprehensive prefetch pipeline with a real-time progress indicator: mutation retrieval, structure selection and download, mutation-to-structure mapping, density calculation, domain annotation, drug interaction queries, clinical trial search, survival data retrieval, and co-occurrence computation. All results are cached in session state, enabling instantaneous tab switching without redundant API calls.

A PDF report export function generates a multi-page summary document containing mutation density charts, hotspot tables, drug interaction listings, survival statistics, and domain annotations, suitable for inclusion in lab meeting presentations or grant applications.

## 4. Comparison with Existing Tools

As shown in Table 1, OncoMORPHIA is the only platform that combines 3D structural visualization with mutation density heatmaps, drug-target overlay, survival analysis, mutational signatures, PPI networks, and AI-powered interpretation in a single interface. While individual features are available across existing tools, the integration eliminates the need for researchers to navigate between multiple platforms and manually cross-reference results.

**Table 1.** Feature comparison of OncoMORPHIA with existing cancer genomics visualization platforms. ✓indicates full support; — indicates feature not available.

| Feature | OncoMORPHIA | cBioPortal | COSMIC | steinPaint | CSB PDB |
| --- | --- | --- | --- | --- | --- |
| 3D structure viewer | ✓ | — | — | — | ✓ |
| Mutation heatmap on structure | ✓ | — | — | — | — |
| Interactive lollipop plot | ✓ | ✓ | ✓ | ✓ | — |
| ClinVar integration | ✓ | Partial | ✓ | — | — |
| Drug-target overlay | ✓ | — | — | — | — |
| Survival analysis | ✓ | ✓ | — | — | — |
| Mutational signatures | ✓ | ✓ | ✓ | — | — |
| PPI network | ✓ | Partial | — | — | — |
| AI interpretation | ✓ | — | — | — | — |
| AlphaFold pLDDT overlay | ✓ | — | — | — | ✓ |
| Pan-cancer tissue breakdown | ✓ | ✓ | ✓ | ✓ | — |
| PDF report export | ✓ | — | — | — | — |
| No installation required | ✓ | ✓ | ✓ | ✓ | ✓ |
| Fully free / no account | ✓ | ✓ | Limited | ✓ | ✓ |

## 5. Example Use Case: TP53 Mutation Analysis

To demonstrate the analytical workflow, we present a complete analysis of TP53, the most frequently mutated gene in human cancer. Upon selecting TP53 and initiating analysis, OncoMORPHIA retrieves 628 mutations from ClinVar (germline classifications) and cBioPortal MSK-IMPACT (somatic). The platform automatically selects PDB entry 5MHC (resolution 1.20 Å, UniProt P04637) as the best-coverage experimental structure, mapping 365 of 628 mutations onto resolved residues, of which 162 carry pathogenic classifications.

The 3D viewer reveals that mutations cluster heavily across the DNA-binding core domain, with nine hotspot residues carrying five or more independent mutation reports. The top hotspots are positions 134 and 208 (6 mutations each) and position 105 (5 mutations). Pathogenicity-coloured spheres immediately distinguish pathogenic (red), likely pathogenic (orange), VUS (yellow), somatic (purple), and benign (green) variants in the structural context. Hovering over any sphere displays the specific HGVS mutation name and classification—for example, residue 164 shows p.Lys164Asn classified as VUS, while residue 136 shows three somatic variants including p.Q136P and p.Q136E.

The interactive lollipop plot provides a complementary linear view, with the UniProt domain track revealing that the highest mutation density falls within the Interaction with ASPP and Transcription activation domains. The Plotly-based hover tooltip at residue 208 shows p.Asp208Tyr, p.Asp208Ala, and p.Asp208His with a “Conflicting” classification— directly flagging this position for further curation.

The Drugs & Trials tab reveals that while no direct drug-gene interactions are catalogued in DGIdb or ChEMBL for TP53 (consistent with p53’s role as a tumour suppressor rather than a conventional druggable target), the platform identifies multiple actively recruiting clinical trials including NCT06659614 (Prostate Tissue BioBank), NCT04367246 (Li-Fraumeni Syndrome/TP53 Biobank), NCT06504199 (Phase 2 combination therapy with Obinutuzumab and Zanubrutinib), and NCT03850574 (Phase 1/2 pharmacokinetic study).

The Survival tab reveals a clear separation between TP53-mutated (n=3,788) and wild-type (n=6,937) patients in the TCGA Pan-Cancer Atlas, with a 35.3% mutation rate across 32 cancer types. Median overall survival is 55 months for mutated patients versus 95 months for wild-type—a 40-month difference visible in the Kaplan-Meier curves with non-overlapping confidence intervals. The AI assistant, when asked about top hotspot residues, provides an immediate context-aware response grounded in the loaded data, listing pos134, pos208, pos105, pos127, and pos158 with their counts.

This complete multi-modal analysis—which would typically require 2–3 hours of manual navigation across ClinVar, cBioPortal, RCSB PDB, DGIdb, ClinicalTrials.gov, and the literature—is completed in OncoMORPHIA in under 90 seconds with a single click.

## 6. Figure Legends

**Figure 1.**
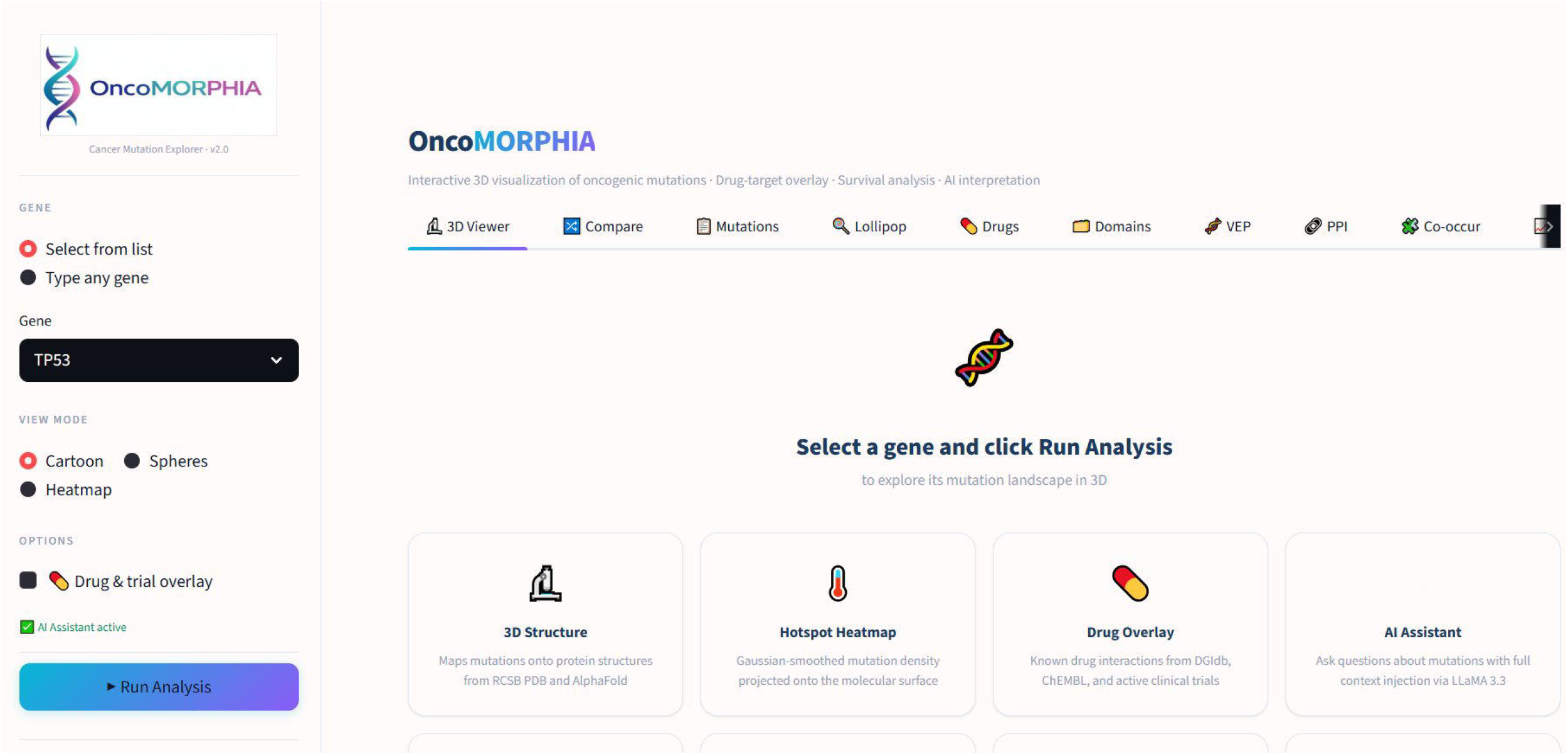
OncoMORPHIA landing page and interface overview. The application presents a clean, white-background interface with the OncoMORPHIA logo and DNA helix branding in the sidebar. Gene selection (dropdown or free text), view mode (Cartoon, Spheres, Heatmap), and analysis options are accessible in the left panel, with a gradient “Run Analysis” button. The main area displays eight feature cards describing the platform’s analytical modules. The tab bar provides access to thirteen analysis tabs: 3D Viewer, Compare, Mutations, Lollipop, Drugs, Domains, VEP, PPI, Co-occur, Survival, Signatures, Pathways, and AI. A green indicator confirms that the AI Assistant is active.

**Figure 2.**
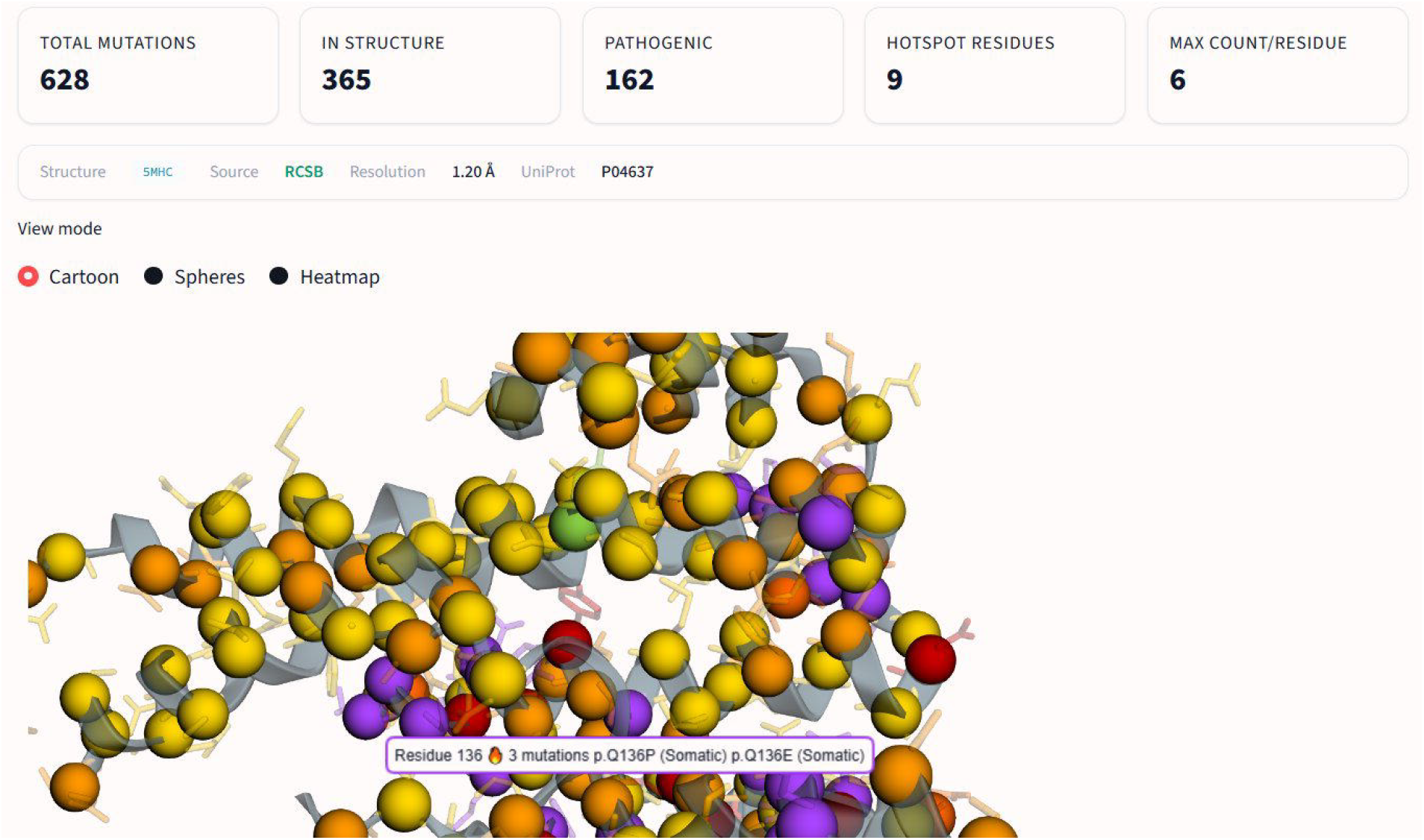
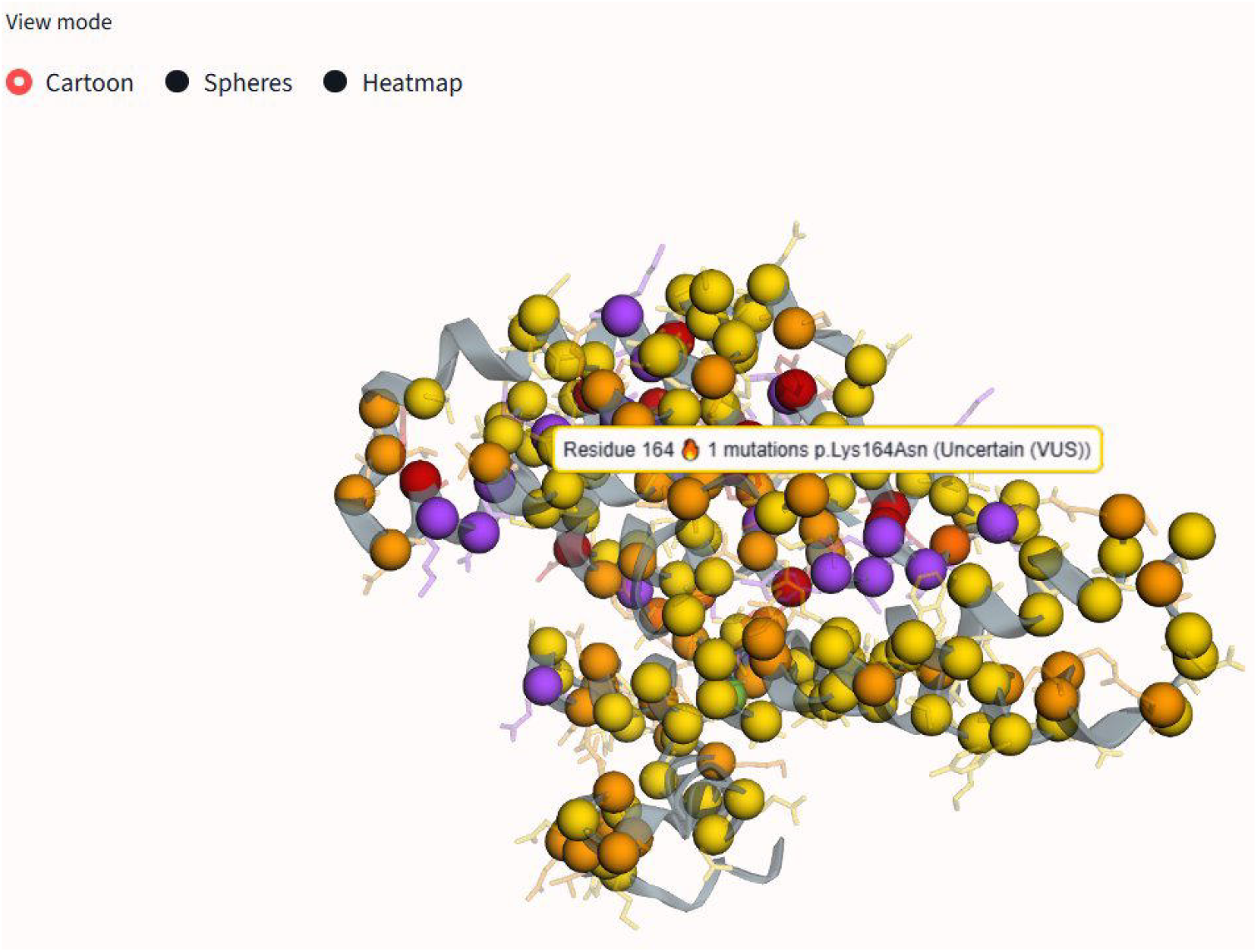
3D protein structure visualization of TP53 mutations. (A) Overview showing summary metrics: 628 total mutations, 365 mapped to structure, 162 pathogenic, 9 hotspot residues, maximum 6 mutations per residue. The structure information bar shows PDB entry 5MHC from RCSB at 1.20 Å resolution (UniProt P04637). The 3D viewer renders the p53 crystal structure in cartoon mode with pathogenicity-coloured mutation spheres: yellow (VUS/uncertain), orange (likely pathogenic), red (pathogenic), and purple (somatic). (B) Close-up view showing interactive hover tooltip at residue 164, displaying “1 mutations p.Lys164Asn (Uncertain (VUS))”. At residue 136, the tooltip shows “3 mutations p.Q136P (Somatic) p.Q136E (Somatic)”, demonstrating the per-residue annotation detail.

**Figure 3.**
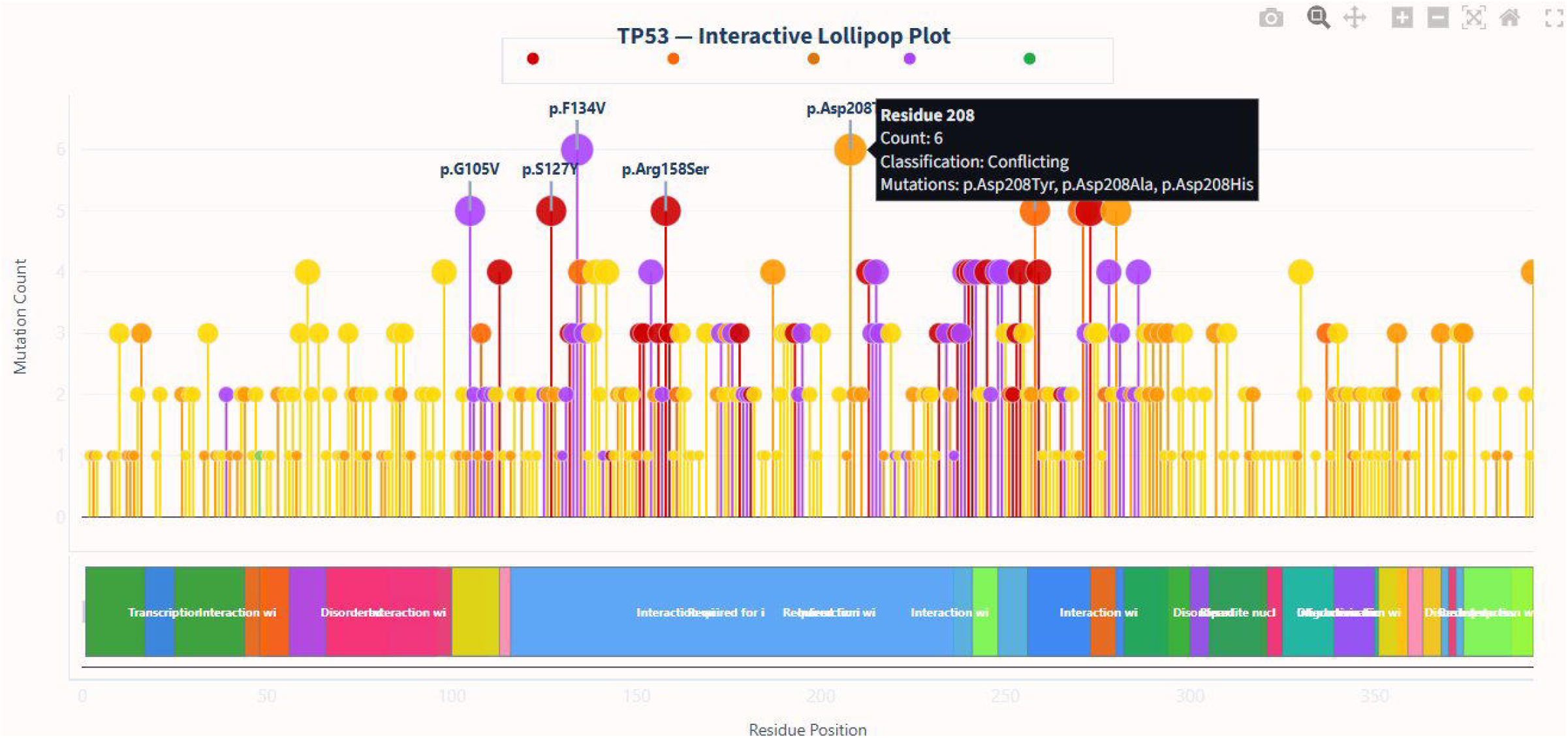
Interactive Plotly-based lollipop plot for TP53. Vertical stems at each mutated residue position carry circles sized by mutation count and coloured by clinical classification: red (pathogenic), orange (likely pathogenic/conflicting), yellow (VUS), purple (somatic), green (benign). Top hotspots are labelled with HGVS notation (p.F134V, p.G105V, p.S127V, p.Arg158Ser, p.Asp208). The hover tooltip at residue 208 shows “Count: 6, Classification: Conflicting, Mutations: p.Asp208Tyr, p.Asp208Ala, p.Asp208His.” The UniProt protein domain track below shows coloured rectangles for annotated domains including Transcription activation, Interaction, and Disordered regions. The Plotly toolbar (top right) provides zoom, pan, and SVG export functionality.

**Figure 4.**
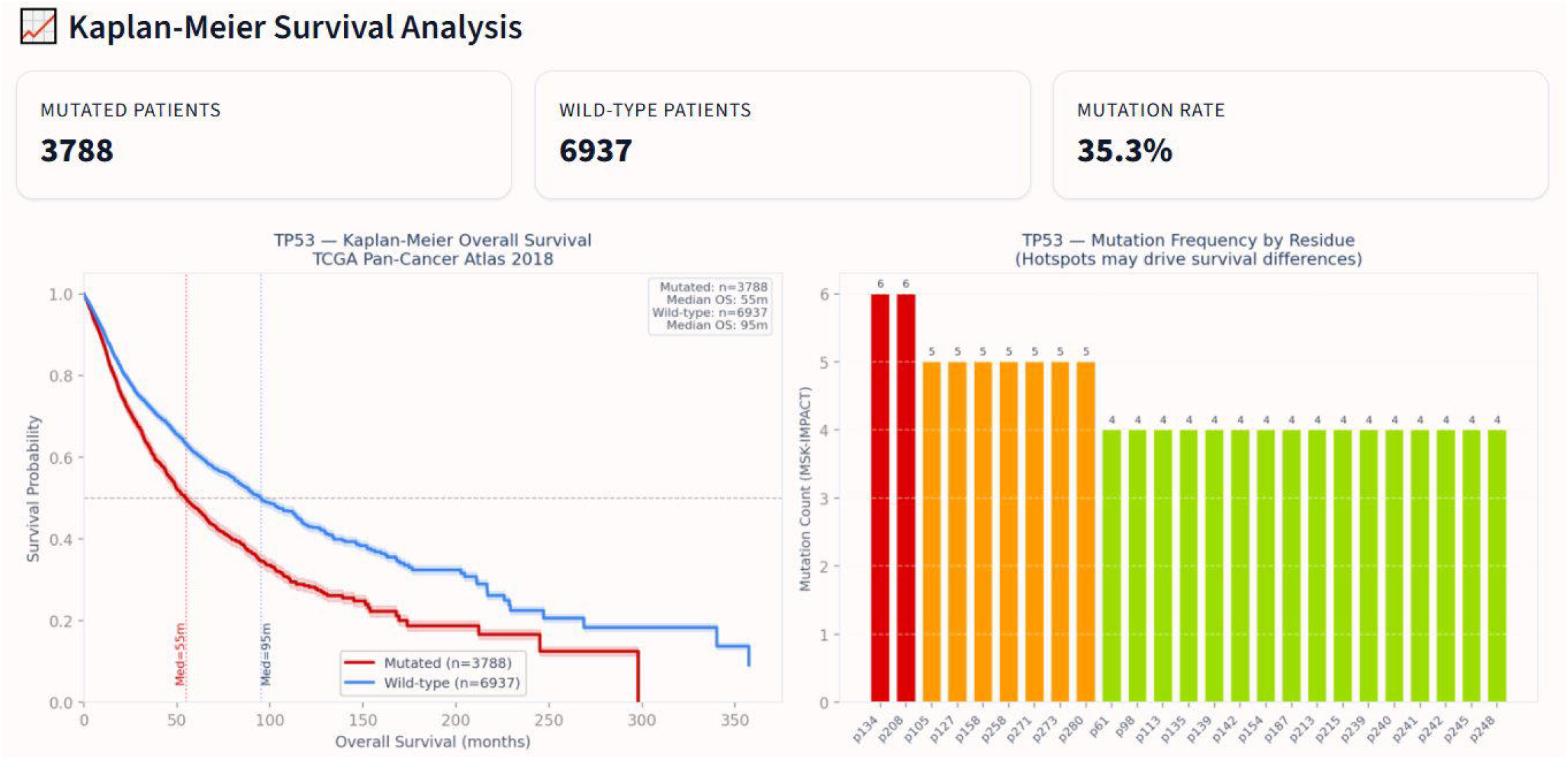
Kaplan-Meier overall survival analysis for TP53. Left panel: Summary metrics showing 3,788 mutated patients, 6,937 wild-type patients, and a 35.3% mutation rate across the TCGA Pan-Cancer Atlas. The Kaplan-Meier curves show clear separation between mutated (red) and wild-type (blue) cohorts, with median overall survival of 55 months for mutated patients versus 95 months for wild-type—a 40-month difference. Dotted vertical lines mark median survival for each group. Shaded bands represent 95% confidence intervals. Right panel: Mutation frequency by residue position from MSK-IMPACT, with bars coloured by a density gradient (red = highest frequency, green = lower), showing the top 25 most frequently mutated positions.

**Figure 5.**
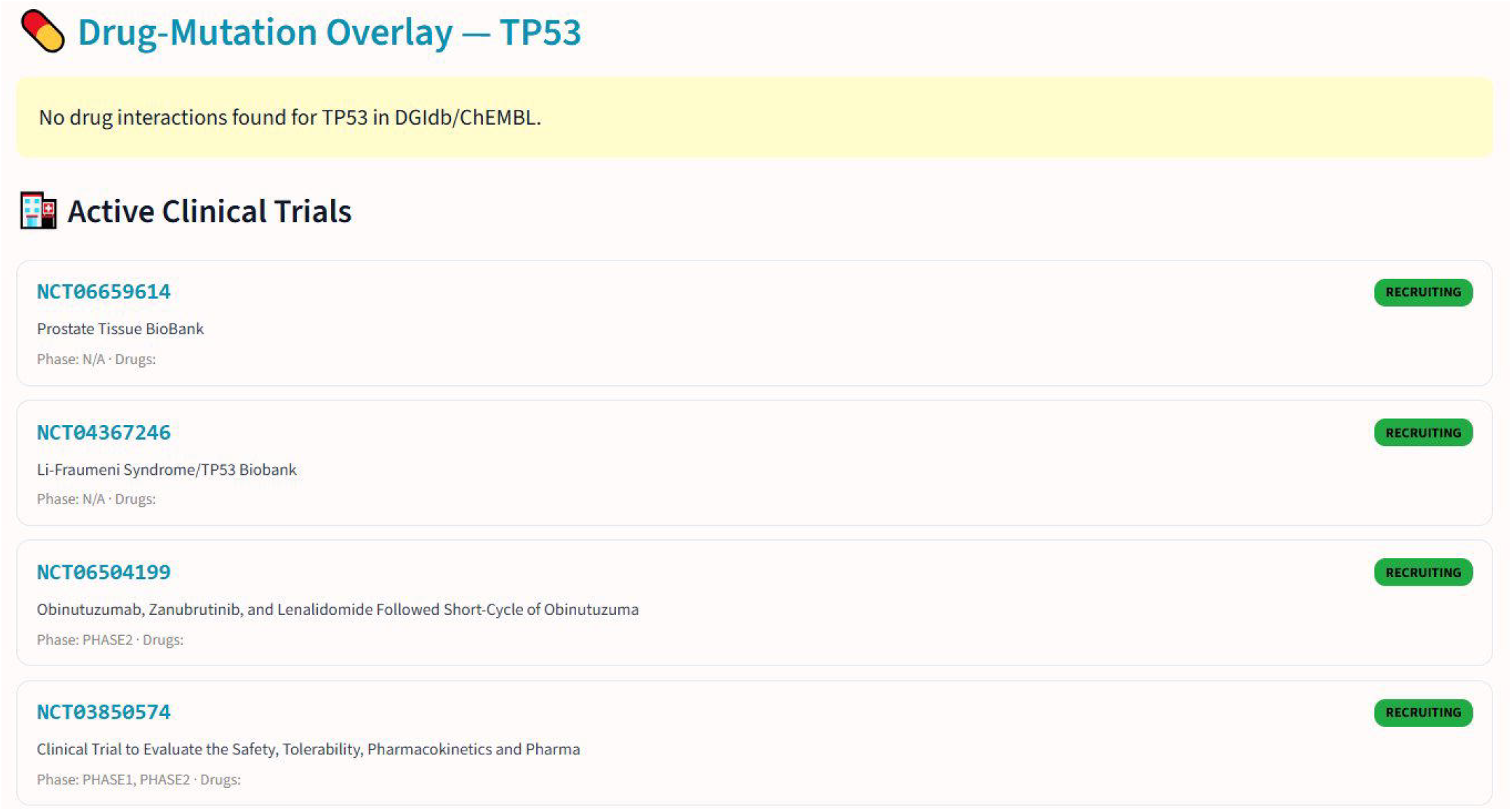
Drug-target interaction and clinical trials panel for TP53. The platform reports that no direct drug-gene interactions are catalogued for TP53 in DGIdb or ChEMBL, consistent with p53’s role as a tumour suppressor. Below, actively recruiting clinical trials are listed with NCT identifiers, trial titles, phase information, and green “RECRUITING” status badges. Trials shown include NCT06659614 (Prostate Tissue BioBank), NCT04367246 (Li-Fraumeni Syndrome/TP53 Biobank), NCT06504199 (Phase 2, Obinutuzumab/Zanubrutinib/Lenalidomide), and NCT03850574 (Phase 1/2, safety and pharmacokinetics). Each NCT identifier links directly to the ClinicalTrials.gov study page.

**Figure 6.**
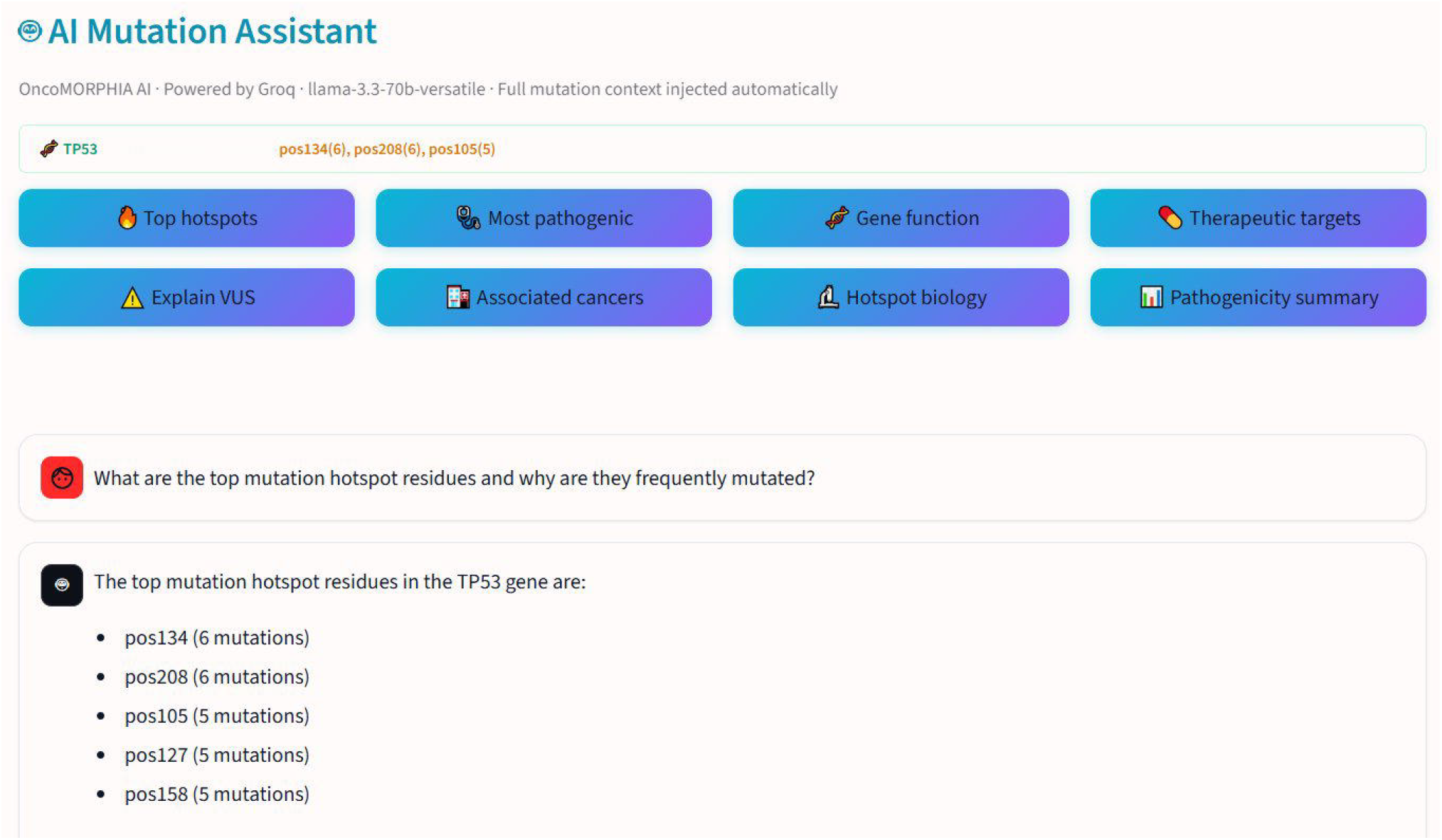
AI-powered mutation interpretation for TP53. The OncoMORPHIA AI assistant (powered by Groq/LLaMA 3.3 70B) displays a context bar showing the loaded gene (TP53) with top hotspots pos134(6), pos208(6), pos105(5). Eight quick-question buttons provide one-click access to common queries. The user asked “What are the top mutation hotspot residues and why are they frequently mutated?” and the assistant responds with a data-grounded answer listing pos134 (6 mutations), pos208 (6 mutations), pos105 (5 mutations), pos127 (5 mutations), and pos158 (5 mutations)— directly reflecting the loaded mutation dataset rather than generic training knowledge.

## 7. Discussion

OncoMORPHIA addresses a critical gap in the cancer genomics toolkit: the lack of an integrated, freely accessible platform that combines structural visualization with comprehensive functional annotation. By automating the retrieval and cross-referencing of data from ten public databases, OncoMORPHIA reduces a multi-hour manual workflow to a single-click analysis.

A key technical contribution is the handling of the January 2024 ClinVar API restructuring. Many bioinformatics tools that query ClinVar programmatically were broken by the transition from the single clinical_significance field to the tripartite germline/oncogenicity/clinical_impact classification system. OncoMORPHIA’s parser probes all four fields (including the legacy format) and applies longest-match-first substring resolution to correctly classify variants, ensuring forward compatibility with future ClinVar schema changes.

The TCGA Pan-Cancer survival analysis required solving a non-obvious data integration challenge. The TCGA Pan-Cancer Atlas is not a single study on cBioPortal but is distributed across 32 individual cancer-type studies (e.g., brca_tcga_pan_can_atlas_2018, skcm_tcga_pan_can_atlas_2018). OncoMORPHIA uses the POST /mutations/fetch endpoint with all 32 molecular profile identifiers to query mutations across the full atlas in a single API call, then aggregates patient-level survival attributes per study.

The inclusion of AI-powered interpretation represents a deliberate design decision. While AI-generated content should never substitute for expert clinical judgment, the context-aware assistant provides a natural language interface for rapidly exploring mutation data, generating hypotheses, and drafting summaries. The full injection of loaded mutation data into the system prompt ensures that responses are specific to the user’s analysis rather than generic.

### 7.1 Limitations

OncoMORPHIA has several limitations that should be acknowledged. First, the survival analysis aggregates all cancer types, which may confound gene-specific effects with cancer-type composition differences. Second, the Kaplan-Meier implementation does not include a log-rank test statistic, though this is planned for a future release. Third, the VEP predictions depend on Ensembl’s ability to resolve protein-level HGVS notation to transcript coordinates, which fails for some non-canonical transcripts. Fourth, the tool currently focuses on missense mutations; frameshift, nonsense, and splice-site variants are included in some modules (co-occurrence, survival) but not in the 3D structural mapping. Fifth, as with any tool that aggregates data from multiple sources, the accuracy of results depends on the currency and completeness of the underlying databases.

### 7.2 Future Directions

Planned enhancements include: implementation of the log-rank test for survival curve comparison; cancer-type-specific survival filtering; integration of evolutionary conservation scores (PhyloP, GERP) for VUS interpretation; batch analysis mode for processing gene lists; and an institutional deployment option with authentication and custom gene panels for clinical genomics laboratories.

## 8. Conclusions

OncoMORPHIA provides a comprehensive, freely accessible web platform for exploring cancer mutations in their structural, clinical, pharmacological, and evolutionary context. By integrating ten public databases into a single analytical environment with interactive 3D visualization, AI-powered interpretation, and one-click report generation, OncoMORPHIA significantly lowers the barrier to entry for cancer mutation analysis and accelerates the translation of genomic data into biological insights.

## Data Availability

OncoMORPHIA is freely available at https://oncomorphia.com. All data displayed in the application is retrieved in real time from public databases (ClinVar, cBioPortal, RCSB PDB, AlphaFold, DGIdb, ChEMBL, ClinicalTrials.gov, STRING DB, UniProt, Ensembl VEP). No proprietary data is used. Source code is available upon request from the corresponding author.

## Acknowledgments

The author acknowledges the developers and maintainers of the public databases and APIs that make OncoMORPHIA possible: NCBI ClinVar, cBioPortal for Cancer Genomics, RCSB Protein Data Bank, the AlphaFold Protein Structure Database, DGIdb, ChEMBL, ClinicalTrials.gov, STRING DB, UniProt, and Ensembl. The author thanks the Streamlit, py3Dmol, Plotly, and BioPython open-source communities.

